# Statistical Colocalization of Genetic Risk Variants for Related Autoimmune Diseases in the Context of Common Controls

**DOI:** 10.1101/020651

**Authors:** Mary D Fortune, Hui Guo, Oliver Burren, Ellen Schofield, Neil M Walker, Maria Ban, Stephen J Sawcer, John Bowes, Jane Worthington, Anne Barton, Steve Eyre, John A Todd, Chris Wallace

## Abstract

Identifying whether potential causal variants for related diseases are shared can increase understanding of the shared etiology between diseases. Colocalization methods are designed to disentangle shared and distinct causal variants in regions where two diseases show association, but existing methods are limited by assuming independent datasets. We extended existing methods to allow for the shared control design common in GWAS and applied them to four autoimmune diseases: type 1 diabetes (T1D); rheumatoid arthritis; celiac disease (CEL) and multiple sclerosis (MS). Ninety regions associated with at least one disease. In 22 regions (24%), we identify association to precisely one of our four diseases and can find no published association of any other disease to the same region; some of these may reflect effects mediated by the target of immune attack. Thirty-three regions (37%) were associated with two or more, but in 14 of these there was evidence that causal variants differed between diseases. By leveraging information across datasets, we identified novel disease associations to 12 regions previously associated with one or more of the other three autoimmune disorders. For instance, we link the CEL-associated *FASLG* region to T1D and identify a single SNP, rs78037977, as a likely causal variant. We also highlight several particularly complex association patterns, including the *CD28-CTLA4-ICOS* region, in which it appears that three distinct causal variants associate with three diseases in three different patterns. Our results underscore the complexity in genetic variation underlying related but distinct autoimmune diseases and help to approach its dissection.

## Introduction

Overlaps of genetic association to different diseases have been widely observed, and are thought to reflect shared etiology between diseases.^1^ However, showing that a variant is associated with two traits does not demonstrate that it is causal for both: this may be due to distinct variants in linkage disequilibrium.^2^ Colocalization analyzes are used to study whether potential causal variants are shared by combining information across multiple single-nucleotide polymorphisms (SNPs) in a region. The proportional approach^3^ tests a null hypothesis of proportionality under which, if causal variants are shared, we expect to see that the effects of any set of SNPs on the two diseases are proportional to each other. A weakness of this approach is interpretation. Failure to reject the null hypothesis does not only imply colocalization, but could also be caused by neither disease being associated, or by insufficient power owing to too few samples analyses and/or an incomplete genetic map^4^ (Supplementary Fig. 1). We have no way of measuring how likely colocalization is. A strength is that no assumptions are made about the number of causal variants: the null hypothesis corresponds to complete sharing across all causal variants. An alternative is to use a Bayesian framework,^5^ to generate posterior probabilities for colocalization and distinct causal variants, as competing hypotheses. However, a weakness of this approach, as currently developed, is that it assumes only a single causal variant for each trait within any region.

Existing colocalization methods require that genetic association with the two traits of interest has been tested in distinct samples. However, this requirement restricts the applicability of the approach to related diseases since each set of case samples must have a corresponding distinct set of control samples, enabling a logistic binomial model to be used independently upon each disease. In contrast, many studies use a common set of controls for different diseases to increase efficiency. Here, we extend both colocalization methods to allow for the use of multinomial logistic regression, the natural model for shared controls.

**Figure 1:**
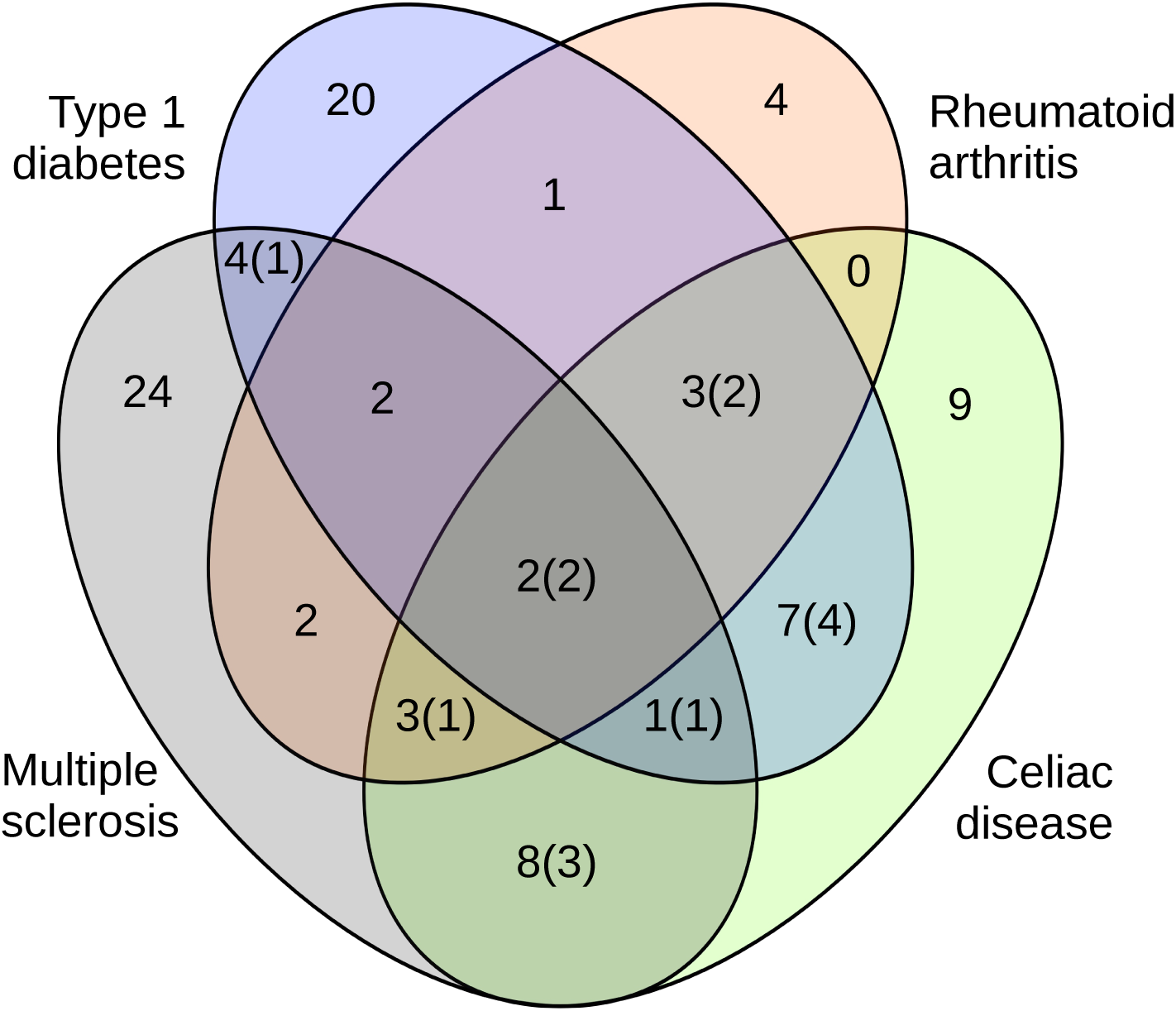
A Venn diagram showing summary of disease assignments to 90 regions which showed association to at least one disease, based upon the results of the Bayesian analysis. In cases where assignment was uncertain, the assignment most supported by the posterior probabilities was used. The numbers in brackets correspond to how many of these regions show evidence of distinct causal variants. Thirty six regions analyzed did not demonstrate association to any disease within our available data, and so are not included in this figure.

Previous studies have identified many regions associated with multiple autoimmune or autoinflammatory diseases, including type 1 diabetes (T1D) and celiac disease (CEL).^6,1^ Such multidisease association led to the development of the ImmunoChip,^7^ a custom genotyping chip with 196,000 SNPs designed to densely cover 186 regions known to associate with at least one immune disease on the basis of GWAS p-value < 10^−8^. The ImmunoChip consortium used a common control set. We applied our extended methods to ImmunoChip raw genotyping data for a total of 36,030 samples, including one set of controls and four disease cohorts, in order to better understand the extent of shared genetic etiology in these diseases.

## Results

The Bayesian method derives the posterior support for each of five hypotheses describing the possible association of the region with both diseases. Of greatest interest are:

ℍ_3_: Both diseases are associated with the region, with different causal variants.

ℍ_4_: Both diseases are associated with the region, and share a single causal variant.

Association with both traits corresponds to ℍ_3_ or ℍ_4_; colocalization corresponds to ℍ_4_. This method requires specification of prior probabilities for each hypothesis. We calibrated priors to match our expectations that about 50% of regions associated with two immune-mediated diseases correspond to a shared causal variant (Supplementary Fig. 2), which is close to the proportion found in a manually curated summary of association to six immune-mediated diseases^8^ (58%). For rheumatoid arthritis (RA)^9^ and multiple sclerosis (MS),^10^ for which only UK subsets of international cohorts were analyzed, we modified priors in regions with published associations to reflect this additional information from the published papers. Where a region was annotated in ImmunoBase as associated with RA or MS, we shrunk our priors for hypotheses corresponding to no association for the disease close towards 0, and increased our priors for the remaining hypotheses (Supplementary Methods).

**Figure 2:**
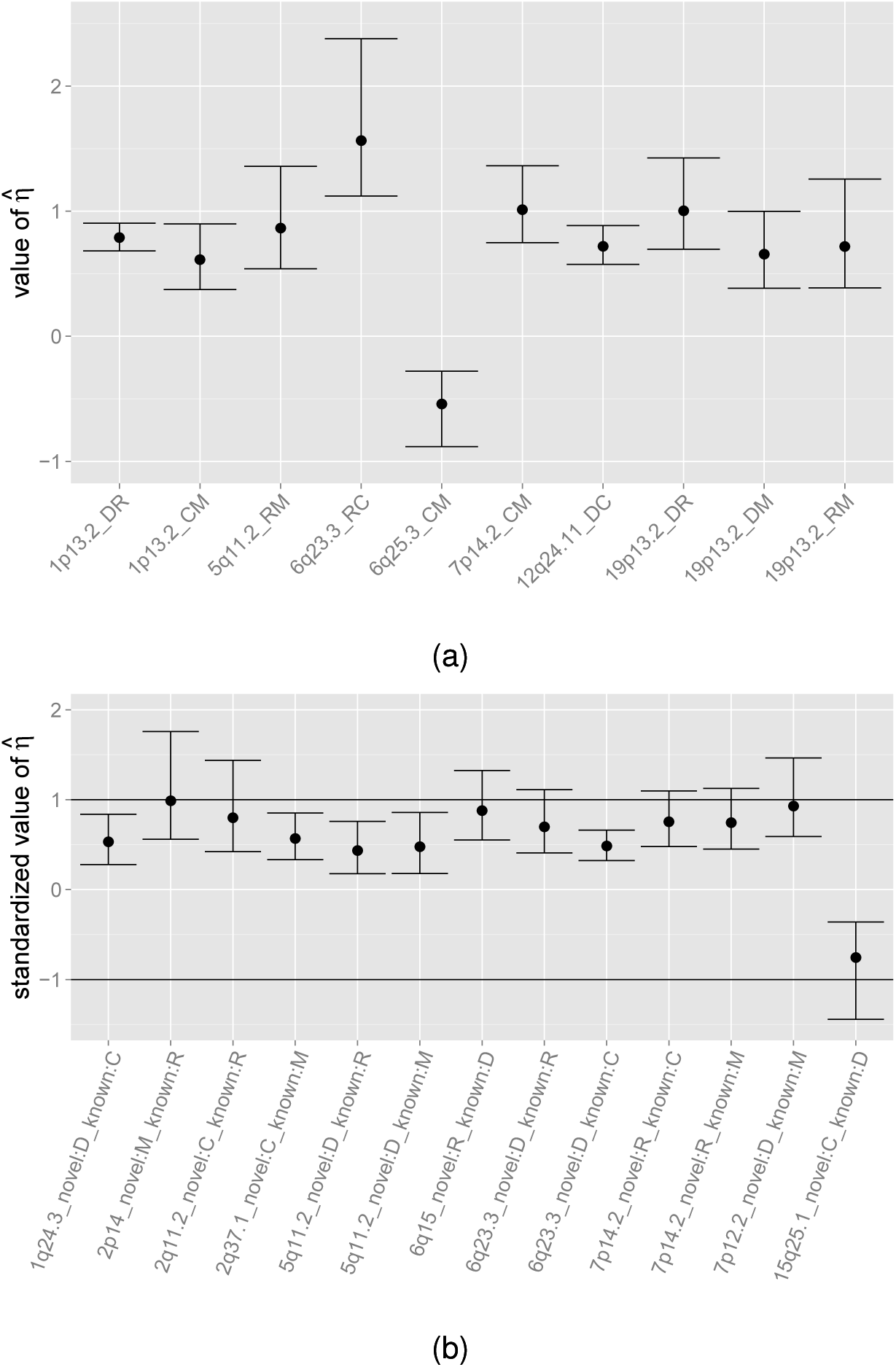
The distribution of 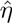, the estimated proportionality coefficient together with its 95% confidence interval. In the case of colocalization, *η* is the ratio of the effects the region exerts upon the two traits. *η* > 1 corresponds to a stronger effect in Trait 2 than Trait 1. We estimate *η* by 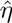. Labels on the x-axis give the traits and regions analyzed; D for T1D, R for RA, C for CEL and M for MS. Note that in some regions, the conditional analysis supports the existance of multiple associated variants: if none of these overlap, then we consider the region to have separate SNP effects. (a) Regions with strong evidence of colocalization (ℙ (*H*4) > 0.9). As we would expect, 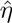 is distributed about 1, which corresponds to the regions having equal effects on each trait. Note that 6q25.3, containing the candidate causal gene *TAGAP*, has 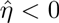, indicating opposite effects on the two diseases. Trait 1 is listed first, and trait 2 second. (b) Regions with novel evidence of disease association, in which we believe there to be colocalisation present between the novel association and at least one of the existing associations. Regions have been ordered such that 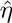 estimates the effect size for the novel trait divided by the effect size for the known association. Labels give the novel association being given first. It can be seen that the effect size tends to be smaller in the new disease.

One hundred and twenty six ImmunoChip regions assigned to at least one of the diseases (based upon knowledge when the chip was designed or identified in subsequent papers and curated in ImmunoBase, http://www.immunobase.org, accessed 12/11/13) were analyzed using both approaches for all six pairwise comparisons of the four diseases. The Bayesian approach assumes a single causal variant per trait in any region. To allow for multiple causal variants, we used a stepwise method. In the overwhelming majority of cases (740 of 756 pairwise comparisons, or 98%), the data were consistent with at most one causal variant per trait in the 126 regions analyzed. In the remaining 16 pairwise comparisons from 8 regions, we use a stepwise method to allow for multiple causal variants. Ninety of the 126 regions (71%) showed association with at least one disease: in 33 regions, the association was shared between at least two diseases (Fig. 1). Complete results are given in Supplementary Table 1, Supplementary Table 2 and Supplementary Table 3). For fifty-seven regions, the greatest support was for association with precisely one of the four diseases: in 21 cases, we know of no other immune-mediated diseases that have reported association to these regions and therefore hypothesize these may be disease specific among autoimmune diseases (Table 1).

**Table 1:**
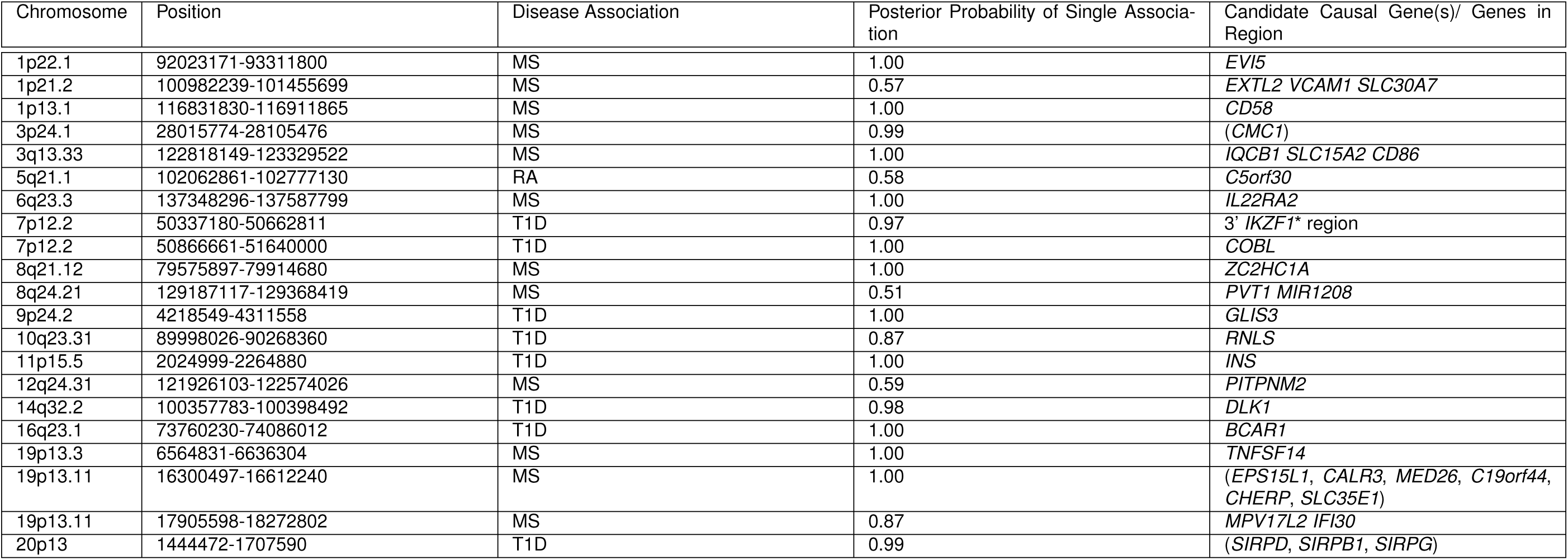
Twenty-one regions which are most likely disease specific under our analysis and for which we know of no other immune-mediated diseases (from the 15 diseases curated in ImmunoBase) that have reported association to these regions (as curated in ImmunoBase, accessed July 9th 2014, and NIHR GWAS catalog, accessed 07/10/2014). Regions required posterior probability of single disease association > 0.5 in at least one pairwise analysis (SNP coverage varies between analyses) and posterior probability of association to any other disorder < 0.2. Candidate causal genes are given. In the case where no candidate causal genes are known, we have given, in brackets, the genes in and around the region. *There are two ImmunoChip regions which overlap *IKZF1* and are separated by a recombination hotspot. The region towards the 5’ end has colocalizing associations with MS and T1D while the region towards the 3’ end appears specific to T1D, as shown in Supplementary Figure 7. Note we provide coordinates of the region, and not an index SNP as is conventional in gwas studies because the method synthesises information across the whole region and does not, in most cases, highlight a single SNP responsible for the association.

**Table 2:**
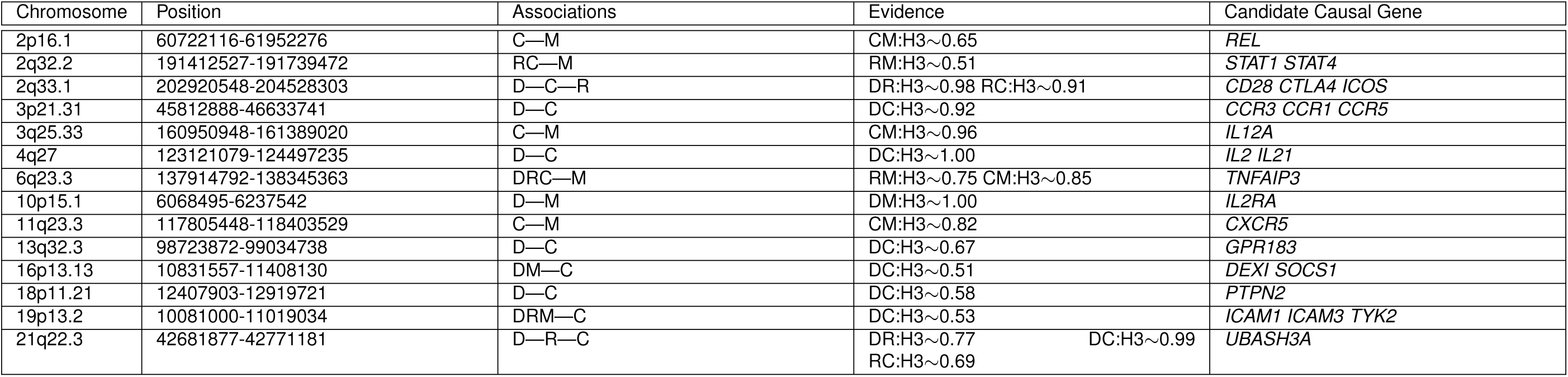
Fourteen regions showing evidence of separate SNP effects (ℙ(*H*3) > 0.5). D corresponds to T1D, R to RA, C to CEL and M to MS. Candidate causal genes are as associated across all curated diseases by ImmunoBase. Distinct signals are indicated by ‘— ’. Many of these regions are associated with other diseases (see ImmunoBase). For instance, the 2q32.2 region is additionally associated with Ulcerative Colitis, Crohn’s Disease, Primary Biliary Cirrhosis, Systemic Lupus Erythematosus and Juvenile Idiopathic Arthritis. The 6q23.3 region is additionally associated with Ulcerative Colitis, Systemic Lupus Erythematosus and Psoriasis. Note that in some regions, eg 10p15.1, the conditional analysis supports the existence of multiple associated variants: if none of these overlap, then we consider the region to have separate SNP effects. Note we provide coordinates of the region, and not an index SNP as is conventional in gwas studies because the method synthesises information across the whole region and does not, in most cases, highlight a single SNP responsible for the association.

**Table 3:**
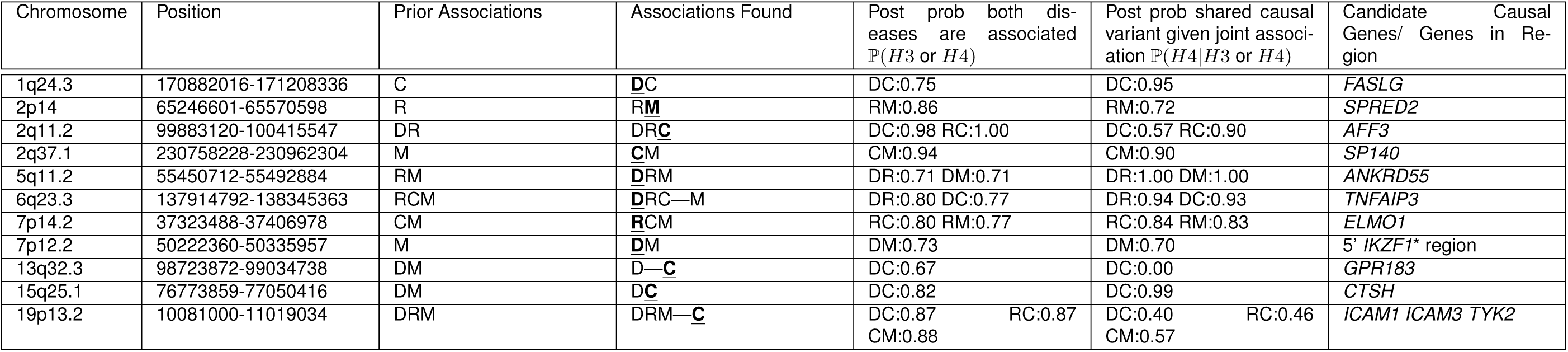
Eleven regions showing strong evidence of novel association (P(*H*3 or *H*4) > 0.5) for an analysis involving a previously non-associated trait. D corresponds to T1D, R to RA, C to CEL and M to MS. Novel associations are underlined and denoted by bold font. Candidate causal genes are as associated across all curated diseases by ImmunoBase. Note that in the case of *TNFAIP3*, there is strong evidence that MS is caused by a distinct causal variant compared to the other traits. Distinct signals are separated by a ‘— ’. Since we only have a subset of the genotype data, not all of the prior (previously published) associations are seen. *An association of T1D in a region 3’ of *IKZF1*, for which it is hypothesised that *IKZF1* is the candidate causal gene is already known^28^ (see Table 1). The novel association we report here is in a region 5’ of *IKZF1*, and independent of the established association. Note we provide coordinates of the region, and not an index SNP as is conventional in gwas studies because the method synthesises information across the whole region and does not, in most cases, highlight a single SNP responsible for the association.

In the Bayesian approach, when the posterior probability of a hypothesis is close to 0.5, assignment cannot be made with confidence to any single hypothesis. However, in the 30 instances in which both diseases showed very strong evidence of association (ℙ(ℍ_3_ or ℍ_4_) > 0.9), the Bayesian and proportional approaches produced consistent results. For these 30 cases, the proportional null was rejected only in cases in which the Bayesian analysis favored H3, and not rejected in cases where H4 was favored. Focusing on these, the data strongly supported that the same causal variants underlie all diseases in ten cases, while seven showed strong evidence for distinct variants, suggesting that just under half, 42%, of overlapping association signals reflect distinct causal variants.

For colocalized disease regions, the two diseases generally have consistent directions of effect (Fig 2) with the exception of the 6q25.3 region containing candidate gene *TAGAP*, which is associated in our analysis with CEL and MS only: the risk allele for CEL is protective for MS and vice versa (Supplementary Fig. 3). This opposing effect of *TAGAP* alleles has been previously described for T1D and CEL,^6^ although the region did not provide sufficient evidence for association with T1D in the data available to us. A similar effect for the 2q12.1 region containing candidate gene *IL18RAP* has been reported.^6^ However, later data^11^ have not offered support for T1D association to 2q12.1, and, in our analysis, the posterior support is concentrated on CEL association alone.

**Figure 3:**
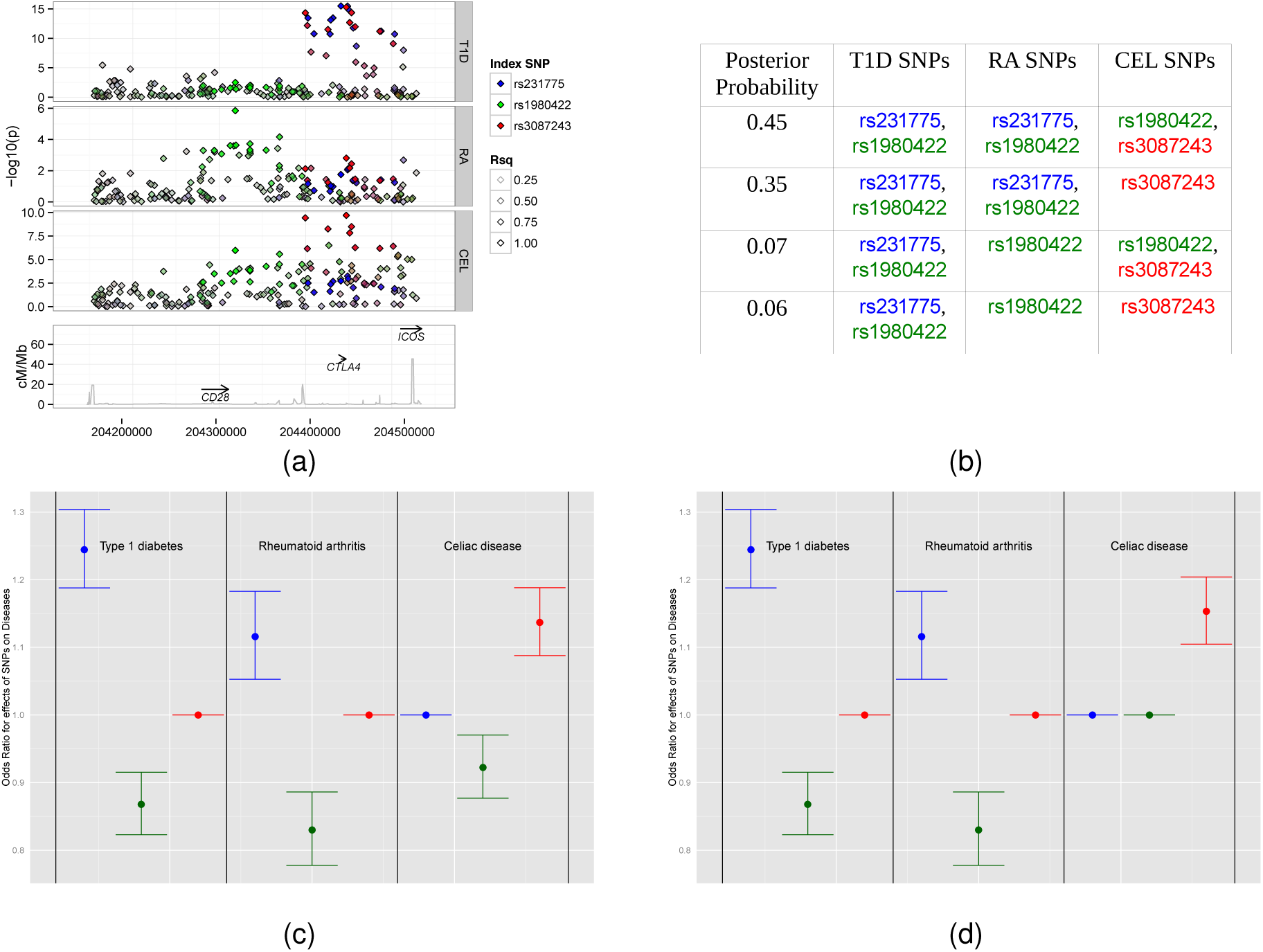
(a) A Manhattan plot of the 2q33.1 region containing the candidate gene *CTLA4*. Three potential causal variants are partially shared between T1D, RA and CEL; the blue signal corresponds to the tag rs231775, the green to rs1980422 and the red to rs3087243. All other SNPs are colored according to their linkage disequilibrium with these three SNPs. SNPs rs231775 and rs3087243 have *r*^2^ = 0.50; all other pairwise *r*^2^ < 1. (b) Each possible model involving these three SNPs was tested; the four models with highest posterior probabilities, which together encompass over 90% of the total posterior probability, are shown. (c) Effect size estimates (including 95% confidence intervals) of each SNP for each disease for the most likely model. (d) Effect size estimates (including 95% confidence intervals) of each SNP for each disease for the second most likely model.

Patterns of association with multiple diseases can be complex. In the 2q33 region containing established candidate gene *CTLA4*, as well as the equally strong functional candidate genes, *CD28* and *ICOS*, three potential causal variants appear to be partially shared between T1D, RA and CEL. The strongest association with T1D is at rs3087243 (which has previously been called CT60), while the strongest association with CEL is with rs231775 (which alters the amino acid at position 17 of CTLA-4, Ala17Thr, and has previously been called CT42). The two SNPs have *r*^2^ = 0.5, and haplotype analysis has previously suggested CT60 and not CT42 is causal for Graves’ disease.^12^ For RA, the strongest single SNP signal is at rs1980422, which is not in LD with either CT42 or CT60 (*r*^2^ < 0.1). We fit each of the 512 possible multinomial models involving these three SNPs for the three diseases. Assuming each model to be equally likely *a priori*, the model with highest posterior probability has rs1980422/rs3087243 (CT60) signals for CEL and rs231775 (CT42)/rs1980422 for both T1D and RA, although while rs231775 (CT42) is the strongest effect for T1D, rs1980422 is strongest for RA (Fig. 3). These results emphasize the potential complexity that can arise in regions of multiple association signals, and motivate the extension of the colocalisation approach developed here to allow model search strategies which does not require stepwise assumptions.

Two regions were associated with all four diseases (Fig. 1). One was the 6q23.3 region containing candidate gene *TNFAIP3*, known to be associated with RA and CEL. There has been some published evidence that T1D is associated with this region,^13^ although not at genome-wide significant levels. Our results identify a T1D signal, colocalized with that for RA and CEL, suggesting a single shared causal variant affecting the three diseases. There is also evidence of MS association, driven by a distinct causal variant (in the CEL-MS analysis, ℙ(ℍ_3_) = 0.83, Supplementary Fig. 4).

The second region was 19p13.2, known to be associated with T1D, RA and MS, containing the strong functional candidate gene *TYK2*, although immune adhesion genes *ICAM1* and *ICAM3* are also good candidate genes. Our analysis supports these associations, with a posterior probability of colocalization approaching 1. We also find evidence for a novel CEL association. In each of the pairwise analyzes involving CEL, the probability of both diseases being associated *∼* 0.88, although this could be a distinct signal: we have ℙ(*H*4*|H*3 or *H*4) *∼* 0.5 (Supplementary Fig. 5). In total, 11 regions showed strong evidence of novel association with ℙ(*H*3 or *H*4) > 0.5 (Table 3).

In regions with colocalising novel associations, effect sizes tended to be smaller in the new disease (Fig. 2). This could indicate that the stronger effect is in the previously known association, or it could be due to Winner’s Curse,^14^ with the previously known associations displaying inflated effect size estimates. In general for colocalized signals, the coefficient of proportionality is centered about 1.

One novel association found was in the chromosome 1q24.3 region, known to be associated with CEL and containing candidate gene *FASLG*. Pathway analysis also produced evidence for a T1D-associated variant here,^15^ although no SNP has reached the genome-wide significance threshold. Our results support a shared causal variant for T1D and CEL (posterior probability 0.71). Our Bayesian approach also enables fine-mapping when dense genotyping data are available, as is the case here. We identified a single likely causal variant lying in a region with strong evidence of predicted regulatory activity, rs78037977 (Supplementary Fig. 6), with a posterior probability of being causal amongst all genotyped variants, given the colocalization hypothesis, of 0.99. Note that rs78037977 was removed from the CEL data in the original analysis^16^ owing to failing a missingness check (the call rate of 99.942% was just below the 99.95% cut-off).

## Discussion

Colocalization methods so far have allowed for the simultaneous analysis of only two traits: a potential weakness when considering more than two diseases, as investigated here. The Bayesian approach could be extended to arbitrarily many traits, at the cost of increased computational complexity and spreading the posterior over an exponentially increasing hypothesis space, potentially making it difficult to draw firm conclusions. Wen et al, in their description of an alternative method for partitioning the association of a single SNP amongst multiple related quantitative traits,^17^ suggest dealing with this complexity by considering only the extremes - a SNP is associated to all traits, exactly one, or none. Such reduction is impractical when analyzing regions, since it does not allow for overlapping but distinct signals. Although we have extended our software to consider three diseases simultaneously, we have chosen for practical reasons to focus on pairwise analyzes with manual curation of the 11 cases (9%) for which more than two diseases showed association.

By analyzing regions known to associate with one disease, we were able to link 12 to additional disorders: in most cases (8/12) the novel disease association was clearly colocalized with a previously known signal, whilst in one case the evidence supported a distinct causal variant for the novel association. In others (3/12) the evidence for colocalization was more equivocal, even with evidence for pairwise association. We also identified 22 regions which appeared associated to only one autoimmune disease. Given the establised influence of sample size on power to detect associations,^18^ and given that many of these regions contain genes linked to immune function, we expect the number of disease specific results to reduce as sample sizes for each disease continue to increase. Indeed, the chromosome 19p13.11, associated with MS in our analysis, has previously been associated with lymphocyte count,^19^ with high LD between the peak MS SNP (rs1870071) and the lymphocyte count SNP (rs11878602, *r*^2^ = 0.99), suggesting an immune mechanism for the association. However, in the case of T1D, two disease-unique regions overlap known type 2 diabetes (T2D) regions. Chromosome 9p24.2, containing the candidate gene *GLIS3*, has been associated with T2D^20^ and fasting glucose^21^ with high LD between the peak SNP for T1D (rs10814914) and these other traits (rs7041847, *r*^2^ > 0.9). *GLIS3* and its causal allele alter disease risk by altering pancreatic beta-cell function, probably by increasing beta-cell apoptosis.^22^ Chromosome 16q23.1, containing the candidate gene *BCAR1*, is associated with T1D in our analysis and T2D,^20^ and the T2D alleles in this region have been associated with reduced beta cell function,^23^ again with high LD between the peak SNPs for T1D (rs8056814) and T2D (rs7202877, *r*^2^ = 0.81). Inspecting the distribution of T2D GWAS p values at the peak SNPs in our T1D associated regions (Supplementary Fig. 7), we note that the peak SNP in the T1D associated region 6q22.32, rs17754780, also shows association to T2D (*p* = 7.9 × 10^−5^) and is in tight LD with peak T2D SNP in the region (rs9385400, *r*^2^ = 0.97). This region has been reported as associated with T2D at genomewide significance in a larger study.^24^ Chromosome 6q22.3 is not uniquely associated to T1D in our analysis because it overlaps an established Crohn’s disease region,^25^ but the lead Crohn’s SNP (rs9491697) is not in LD with the T1D SNP (*r*^2^ = 0.03). This is then likely to be a third shared signal between T1D and T2D. The nearest genes are *MIR588* about which little appears to be known and *CENPW* (centromere protein W) which is a has no obvious functional candidacy. This genetic overlap between T1D and T2D (Supplementary Table 4) emphasizes that T1D results from an interaction between the immune system and beta cells, and it is probable that some of our other apparent disease unique regions will also prove to be specific to the target of autoimmune destruction in MS and RA.

In a standard GWAS analysis, a p-value significance threshold of 5 × 10^−8^ is used in absence of replication data, due to a desire to minimise reporting of false positive results, although a relaxation of this threshold has been suggested.^26^ However, since autoimmune diseases are known to share etiology, conditioning upon association for one autoimmune disease, we should require a less stringent threshold to believe it significant for another. Indeed, whilst the question of whether the ImmunoChip significance threshold should be somewhat relaxed remains,^8^ examination of p-values in the regions in which we observe novel associations (Supplementary Fig. 8) suggests that a threshold between 10^−5^ and 10^−6^ for SNPs that are confirmed index SNPs for another disease might be more appropriate. Given our estimate that 42% of overlapping and genome-wide significant immune-mediated disease signals relate to distinct causal variants, we suggest that physical proximity to a known associated variant in a related disease, and not only LD with it, does appear an appropriate criterion with which to alter interpretation of a small but not genome-wide significance threshold. Variants meeting such thresholds might be prioritised for genotyping in replication samples. We note, also, that the four diseases we studied are all characterized by the presence of autoantibodies. Had we included autoantibody negative diseases we might have found a higher proportion of discordant associations as reported in a previous manual curation of ImmunoChip studies,^8^ given there remains considerable overlap in location of association signals. Although a careful and detailed manual curation of several studies has been conducted,^8^ the ability of colocalization methods to distinguish shared from distinct causal variants allows clearer interpretation of genetic results.

In summary, we have developed a methodology for examining shared genetic etiology between diseases in the case of common control datasets, extending previous work.^2,3^ This enables the discovery of new disease associations and the exploration of complex association patterns. Although these methods have been presented in this paper to analyze autoimmune diseases, the prior is user defined, and could be used to analyze any pair of related diseases.

## Online Methods

### Samples

All samples included in this analysis were gathered in the United Kingdom, and have reported or self declared European ancestry. Detailed summaries of the sample cohorts are given in the ImmunoChip papers for CEL,^16^ RA,^9^ MS^10^ and T1D (personal communication, Steve Rich). For the RA and MS cases, we used the subset of cases from the UK. Sample exclusions were applied as described in each paper, and in total, 6691 T1D, 3870 RA, 7987 CEL, 5112 MS and 12370 control samples were analyzed. SNPs were filtered according to the following criteria: call rate > 0.99; minor allele frequency > 0.01; HardyWeinberg *|Z|* < 5. SNPs which passed these threshold in controls and any specific pair of cases were used for that pairwise analysis.

### Selection of Regions for Analysis

We considered all regions annotated in ImmunoBase (http://www.immunobase.org, accessed on 12/11/13) as associated with at least one of our diseases. Where regions overlapped, we formed the union. Regions containing fewer than ten SNPs or with a SNP density < 1 SNP/kb were excluded. The MHC (chr6:29797978-33606563 hg18) was removed from the analysis, since this region is known to have complex multi-SNP effects. A full list of the 126 regions analyzed, together with our resulting associations, can be found in Supplementary Table 1.

### Colocalization Analysis

Two colocalization methods were applied to each of the 126 regions (see Supplementary Fig.1).

#### Bayesian Approach

The first approach is based upon a Bayesian approach proposed by Giambartolomei et al.^5^ All models in which each trait is caused by at most one variant are considered, and approximate Bayes factors computed for each. Our extension follows the same framework, but, in order to extend this method to the case of a common control, a multinomial model was used. Bayes factors were computed using a Laplace approximation ^27^ as implemented in the R package mlogitBMA (http://cran.r-project.org/web/packages/mlogitBMA/index.html). Each of these models is contained within precisely one of the following sets:

ℍ_0_: No SNP is associated with either trait.

ℍ_1_: There is a SNP associated with trait 1, but no SNP is associated with trait 2.

ℍ_2_: There is a SNP associated with trait 2, but no SNP is associated with trait 1.

ℍ_3_: Both the diseases are associated with the region, with different causal variants.

ℍ_4_: Both the diseases are associated with the region, and share a single causal variants.

By summing the Bayes Factors generated for all models in the set, a posterior possibility can be computed for each of the hypotheses, and hence for colocalization (ℍ_4_). Similarly, the posterior probability of any given model, given a specific hypothesis and equal prior probability of each model, is proportional to the BF for that model. Since a Bayes factor is assigned to each model independently, it is straightforward to calculate the conditional probability of each SNP being causal, given association, as proportional to the Bayes factor for the relevant model.

This approach assumes a single causal variant at any region. We tested this assumption in regions with strong evidence of association (ℙ (ℍ_0_) < 0.1) by performing conditional analysis. Firstly, all plausibly important SNPs were discovered by iteratively conditioning on the most likely set of SNPs to cause the associations seen, until there was no longer strong evidence of additional association. In those cases where multiple SNPs were considered relevant, all but a pair (one potentially causal for the first trait, and one for the second) were conditioned upon, in order to discover the colocalization (or not) of the effects at this pair alone.

#### Proportional Approach

A second method based upon the proportional approach^2,3^ was also used. Phenotypes are modeled using multinomial logistic regression, producing maximum likelihood estimates *b*_1_ and *b*_2_ of regression coefficients *β*_1_ and *β*_2_. Since the samples sizes can be large, the asymptotic normality of maximum likelihood estimators is used to approximate:

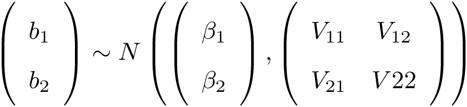

for some variance-covariance matrix **V**.

The method in ^3^ assumes that *b*_1_, *b*_2_ are independent (i.e. *V*_12_ = *V*_21_ = **O**). However in the extension to a common control dataset, we cannot assume this, and proceed with a fully unknown **V**.

The null hypothesis corresponds to the existence of a constant *η* such that 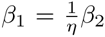. Under this hypothesis, and given *η*,

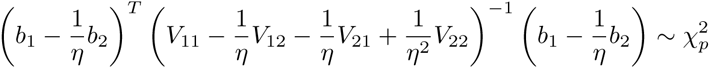

This is used as our test statistic. However, since the value of *η* was unknown, a posterior predictive p-value is generated instead, by integrating the p-values associated with the test statistic over the posterior distribution of *η*. To avoid bias in regression coefficients due to selection of SNPs on the basis of their strength of association, Bayesian model averaging was used to average inference over all plausible two SNP models.

Further details of the colocalization methods can be found in the Supplementary Methods section, and an R package for their implementation is available from https://github.com/mdfortune/colocCommonControl.

### Identification of disease specific regions

To examine evidence for GWAS association with other traits, we took the index SNP with smallest p values in a region, and then identified proxy SNPs based on LD (*r*^2^ *≥* 0.9) using 1000 genomes EUR data. We used this as a query SNP set to examine associations annotated in the NIHR GWAS catalog (http://www.genome.gov/admin/gwascatalog.txtaccessed07/10/2014)

We identified disease specific regions for which: the posterior probability for single SNP association was >0.5; posterior probability of association with any other disease was < 0.2; the region was not annotated as associated with any other autoimmune disease in ImmunoBase; and no proxies for the index SNP were associated with any other autoimmune disease in the NIHR GWAS catalog.

### Type 2 diabetes data

Summary from a T2D GWAS meta analysis^20^ was downloaded from the DIAGRAM website (http://diagram-consortium.org/, accessed 20/10/14).

## Acknowledgments

M.D.F is funded by the Wellcome Trust (099772). C.W and H.G are funded by the Wellcome Trust (089989).

This work was funded by the JDRF (9-2011-253), the Wellcome Trust (091157) and the National Institute for Health Research (NIHR) Cambridge Biomedical Research Centre. This work was funded by the JDRF (9-2011-253), the Wellcome Trust (091157) and the National Institute for Health Research (NIHR) Cambridge Biomedical Research Centre. The Cambridge Institute for Medical Research (CIMR) is in receipt of a Wellcome Trust Strategic Award (100140). ImmunoBase.org is supported by Eli Lilly and Company.

We thank the UK Medical Research Council and Wellcome Trust for funding the collection of DNA for the British 1958 Birth Cohort (MRC grant G0000934, WT grant 068545/Z/02). DNA control samples were prepared and provided by S. Ring, R. Jones, M. Pembrey, W. McArdle, D. Strachan and P. Burton.

This research utilizes resources provided by the Type 1 Diabetes Genetics Consortium, a collaborative clinical study sponsored by the National Institute of Diabetes and Digestive and Kidney Diseases (NIDDK), National Institute of Allergy and Infectious Diseases (NIAID), National Human Genome Research Institute (NHGRI), National Institute of Child Health and Human Development (NICHD), and the JDRF and supported by U01 DK062418.

We are grateful to S. Eyre for use of the rheumatoid arthritis data. Funding was provided by the Arthritis Foundation and the US National Institutes Health.

We are grateful to G. Trynka and D. van Heel for use of the celiac disease data. Funding was provided by the Wellcome Trust, by grants from the celiac disease Consortium and an Innovative Cluster approved by the Netherlands Genomics Initiative, by the Dutch Government (BSIK03009 to C. Wijmenga) and the Netherlands Organisation for Scientific Research (NWO, grant 918.66.620) and by the US National Institutes of Health grant 1R01CA141743 and Fondo de Investigacin Sanitaria grants FIS08/1676 and FIS07/0353.

Multiple sclerosis data was provided by S. Sawcer and the International Multiple Sclerosis Genetics Consortium. Funding was provided by the US National Institutes of Health, the Wellcome Trust, the UK MS Society, the UK Medical Research Council, the US National MS Society, the Cambridge National Institute for Health Research (NIHR) Biomedical Research Centre, DeNDRon, the Bibbi and Niels Jensens Foundation, the Swedish Brain Foundation, the Swedish Research Council, the Knut and Alice Wallenberg Foundation, the Swedish Heart-Lung Foundation, the Foundation for Strategic Research, the Stockholm County Council, Karolinska Institutet, INSERM, Fondation d’Aide pour la Recherche sur la Sclrose en Plaques, Association Franaise contre les Myopathies, Infrastrutures en Biologie Sant et Agronomie (GIS-IBISA), the German Ministry for Education and Research, the German Competence Network MS, Deutsche Forschungsgemeinschaft, Munich Biotec Cluster M4, the Fidelity Biosciences Research Initiative, Research Foundation Flanders, Research Fund KU Leuven, the Belgian Charcot Foundation, Gemeinntzige Hertie Stiftung, University Zurich, the Danish MS Society, the Danish Council for Strategic Research, the Academy of Finland, the Sigrid Juselius Foundation, Helsinki University, the Italian MS Foundation, Fondazione Cariplo, the Italian Ministry of University and Research, the Torino Savings Bank Foundation, the Italian Ministry of Health, the Italian Institute of Experimental Neurology, the MS Association of Oslo, the Norwegian Research Council, the South-Eastern Norwegian Health Authorities, the Australian National Health and Medical Research Council, the Dutch MS Foundation and Kaiser Permanente.

